# Optineurin deficiency induces patchy hair loss but it is not sufficient to cause amyotrophic lateral sclerosis in mice

**DOI:** 10.1101/2021.12.22.473616

**Authors:** Shivranjani C Moharir, Ghanshyam Swarup

## Abstract

Genetic alterations and environmental factors contribute towards pathogenesis of amyotrophic lateral sclerosis (ALS), a neurodegenerative disease. Various types of mutations in the autophagy receptor protein optineurin (coded by *OPTN* gene) including deletions are causatively associated with ALS. To explore the role of OPTN in ALS pathogenesis, we used Optn knockout mice to study the features of ALS. The Optn-deficient mice did not show kyphosis, loss of body weight, weakening of front paw-grip strength or limb muscle strength. However, several Optn-deficient mice showed patchy loss of hair, which increased with age. Our results suggest that optineurin deficiency alone is not sufficient to induce ALS-like symptoms in mice. We suggest that optineurin deficiency may require cooperation with other genetic or environmental factors to cause ALS. Since endoplasmic reticulum (ER) stress plays an important role in ALS pathogenesis, and Optn modulates ER stress response signaling, Optn deficiency may contribute to ALS pathogenesis partly by potentiating ER stress response signaling.

## Introduction

Optineurin is a multifunctional protein involved in many physiological processes like autophagy, NFκB signaling, vesicular trafficking, cell survival, cell division, antiviral signaling, etc. [1, 2]. Certain mutations in the coding region of *OPTN* gene contribute to pathogenesis of glaucoma and amyotrophic lateral sclerosis (ALS), both of which are neurodegenerative diseases [3, 4]. Most of the glaucoma associated mutations in optineurin are likely to be dominant since they are heterozygous missense mutations. The mutations associated with ALS include deletion, truncation and missense mutations. Optineurin mutations leading to ALS pathogenesis possibly function by gain of function as well as loss of function mechanisms [4]. Glaucoma involves degeneration of retinal ganglionic cells whereas ALS involves degeneration of motor neurons of brainstem, primary cortex and spinal cord.

ALS is a fatal disorder characterized by selective and progressive death of motor neurons leading to progressive paralysis, respiratory depression and death. About 90% of the cases of ALS are sporadic whereas the remaining are familial [2]. Mutations in several genes are associated with ALS including *SOD1* (Superoxide Dismutase 1) [5], *TDP43* (TARDP encoding transactive response (TAR) DNA-binding protein-43) [6], *FUS* (fused in sarcoma) [7, 8] and *C9ORF72* (chromosome 9 open reading frame 72) [9, 10]. The exact mechanism underlying the degeneration is still not known but various hypotheses suggest the involvement of oxidative damage, endoplasmic reticulum (ER) stress, transport impairment, toxicity from intracellular aggregates, mitochondrial dysfunction, glutamate mediated excito-toxicity, inflammation, etc. in the pathogenesis of the disease. Several emerging themes suggest that along with the above mentioned molecular and genetic determinants, other exogenous environmental factors also contribute to the pathobiology of ALS [11, 12].

Although loss of function mutations are associated with familial as well as sporadic ALS in humans, it is yet to be established that OPTN deficiency alone is sufficient to cause ALS. Here, we have used *Optn^-/-^* mice to address the role of optineurin in ALS. Our results suggest that Optn knockout (KO) mice do not show symptoms of ALS but these mice show patchy hair loss. These results indicate that Optn deficiency alone is not sufficient to induce ALS in mice and it may require cooperation with other factors (genetic or environmental or both) to induce ALS.

## Materials and methods

### Antibodies

Rabbit polyclonal antibody against Optineurin (Abcam; Cat. No. ab23666; WB 1:1000), mouse monoclonal antibody against Actin (Millipore, Cat. No. MAB1501; WB 1:10000), HRP conjugated anti-mouse IgG (Amersham, NA9310), HRP-linked anti-rabbit IgG (Cell Signaling Technology, 7074) and HRP conjugated anti-rabbit IgG (Amersham, NA934) were used.

### Generation and maintenance of optineurin knockout mice

The Institutional Animal Ethics Committee of Centre for Cellular and Molecular Biology approved all the animal experiments. Optn KO mice were generated by replacing the exon 2 of optineurin gene with β-galactosidase reporter gene which was fused with neomycin resistance gene [13]. This was followed by a polyA sequence [13]. The genotypes of mice were confirmed by performing polymerase chain reaction (PCR) of genomic DNA isolated from the tail of optineurin knockout mice with a pair of primers specific for exon II, as described earlier [14]. The 129S6/SvEvTac cross C57BL/6Ncr ES cell line (G4) [15] was used for generation of Optn KO mice. These ES cells were injected into 3.5 day old blastocysts which were transplanted into uterus of CD1 females. Chimeric males thus obtained were crossed with CD1 females [13]. Thus, the Optn KO mice generated were of mixed background. In order to generate Optn KO mice in C57BL/6 background, the mice were backcrossed for ten generations with C57BL/6 mice.

The mice were maintained in individually ventilated cages in controlled temperature (25°C), humidity and light-dark cycle (12hrs, 6am-6pm) environment. Animals were fed with autoclaved *ad libitum* diet and water. Males and females were maintained in separate cages. Not more than 5 females or 3 males were kept in one cage. The mice were carefully examined at regular intervals. Experimental animals were generated by crossing *Optn^+/-^* male with *Optn^+/-^* female. The progenies followed the Mendelian law, i.e., the ratio of wild type (WT): heterozygous: KO was 1:2:1. WT littermates were used as controls for experiments with Optn KO mice.

### Protein isolation from tissues and western blotting

For protein isolation, the tissues were homogenized in RIPA buffer (50mM Tris pH 7.5, 150mM NaCl, 5mM EDTA, 0.1% SDS, 1% NP-40) supplemented with protease inhibitor cocktail. The sample was incubated on ice for at least 1hour and then centrifuged at 15000 g for 30 minutes at 4°C. The supernatant was taken as RIPA soluble fraction and protein estimation was performed by Bicinchoninic acid method. The protein samples were boiled in 1X SDS-PAGE sample buffer, and western blotting was carried out as described earlier [14].

### Paw grip strength test

In order to test the muscular strength of forelimb of Optn KO and WT mice, paw grip strength test was performed periodically using grip strength meter (MK-380M; Muromachi Kikai Co. Ltd., Tokyo, Japan). The digital values which came in the unit of newton were normalized with the weights of mice.

### Rotating wire test

To assess the muscle strength and co-ordination, we made an apparatus based on the model described earlier [16], consisting of a 50 inches long wire welded into a circle. The circular loop was suspended from a pulley held at a stand. Upon application of downward force, the loop moved freely on the pulley. The mice were positioned with all the four limbs in contact with the loop and the hanging time was recorded thrice with 30 second gap in between each reading. The mean of the three readings of time was taken as the output. To calculate the hanging strength, the time of the hang was multiplied by the weight of the mice as described earlier [16].

### Statistical significance

Statistical significance was calculated using two tailed Student’s T test. p-value less than 0.05 was considered as significant. Error bars represent standard deviation. Hair loss data was analyzed using Chi Squared test with 95% confidence interval.

## Results

The *Optn^-/-^* mice were generated by homologous recombination and these mice were genotyped by PCR as shown in Figure 1A. Absence of full length Optn was confirmed by western blotting (Figure 1B). The C57BL/6 Optn-deficient and wild type littermates were routinely monitored for any obvious phenotypic abnormalities. These mice did not show any obvious ALS-like symptoms such as bending of back (Figure 1C). The phenomenon of bending of spine is known as kyphosis and is observed in TDP43 depleted mice, which develop ALS [17]. However, it was very often observed that some of the Optn KO mice developed patchy hair loss known as alopecia (Figure 1D and E). Wild type mice did not show any such hair loss. The hair loss phenotype of Optn KO mice increased with age, and 9% of 11months old, 16% of 12 months old, 22% of 13 months old and 28% of 14 months old KO mice showed hair loss as compared to 0% corresponding WT control mice (Fig. 1 D).

**Figure 1:**
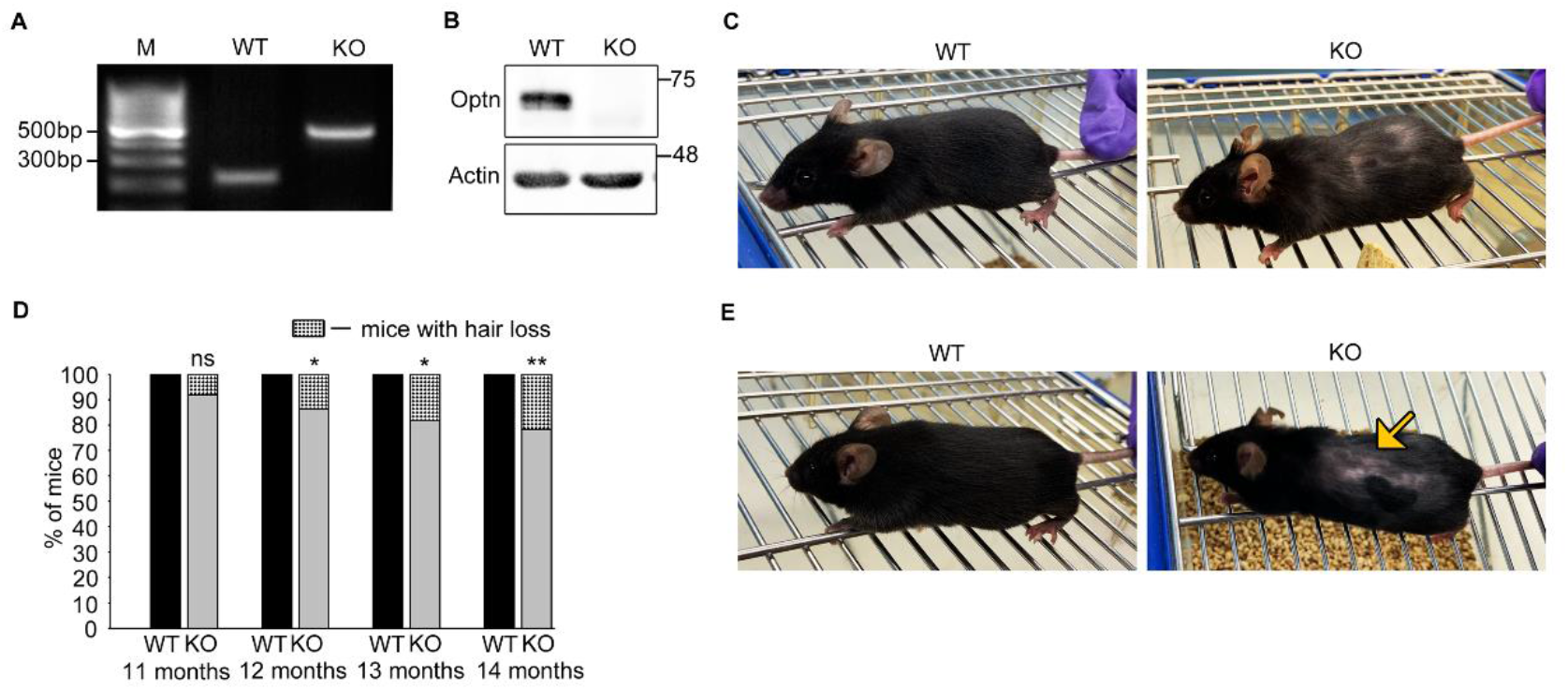
Optineurin knockout mice show patchy hair loss: **(A)** PCRs of tail DNA from optineurin knockout (KO) and wild type (WT) mice. **(B)** Western blot showing absence of optineurin protein (67 kDa) in brainstem of optineurin knockout mice. **(C)** Representative images showing absence of kyphosis in C57BL WT and optineurin knockout mice. **(D)** Bar diagram showing percentage of optineurin knockout mice having hair loss at the age of 11 (15 WT, 23 KO; p=0.247), 12 (27 WT, 32 KO; p=0.033), 13 (25 WT, 41 KO; p= 0.012) and 14 (21 WT, 29 KO; p=0.009) months. *p ≤ 0.05 and **p ≤ 0.01. Significance was calculated using Chi-squared test taking 95% confidence interval. **(E)** Representative images showing hair loss in C57BL optineurin knockout mice.

One of the hallmarks of ALS is loss of motor neurons of spinal cord, brainstem and cortex, which leads to muscle degeneration resulting in weakness of the limbs. Therefore, we measured the paw grip strength, which is a routinely used test to investigate the muscle strength in rodents [18, 19]. We monitored paw grip strength and body weights at 11, 12, 13, 14 and 22 months of age (Figure 2A-H). We observed that there was no significant difference in paw grip strength of Optn KO mice and WT control mice of C57BL/6 background (Figure 2A and D). We also monitored the body weight of the same mice and observed that the Optn KO mice and corresponding WT control mice, did not show any significant difference (Figure 2E and H). We then performed paw grip strength analysis for males and females separately (Figure 2B and C). Optn KO and WT control male mice did not show any significant difference in either the paw grip strength (Figure 2B) or body weight (Figure 2F). Optn KO and WT control female mice also did not show any significant difference in either the paw grip strength (Figure 2C) or body weight (Figure 2G).

**Figure 2:**
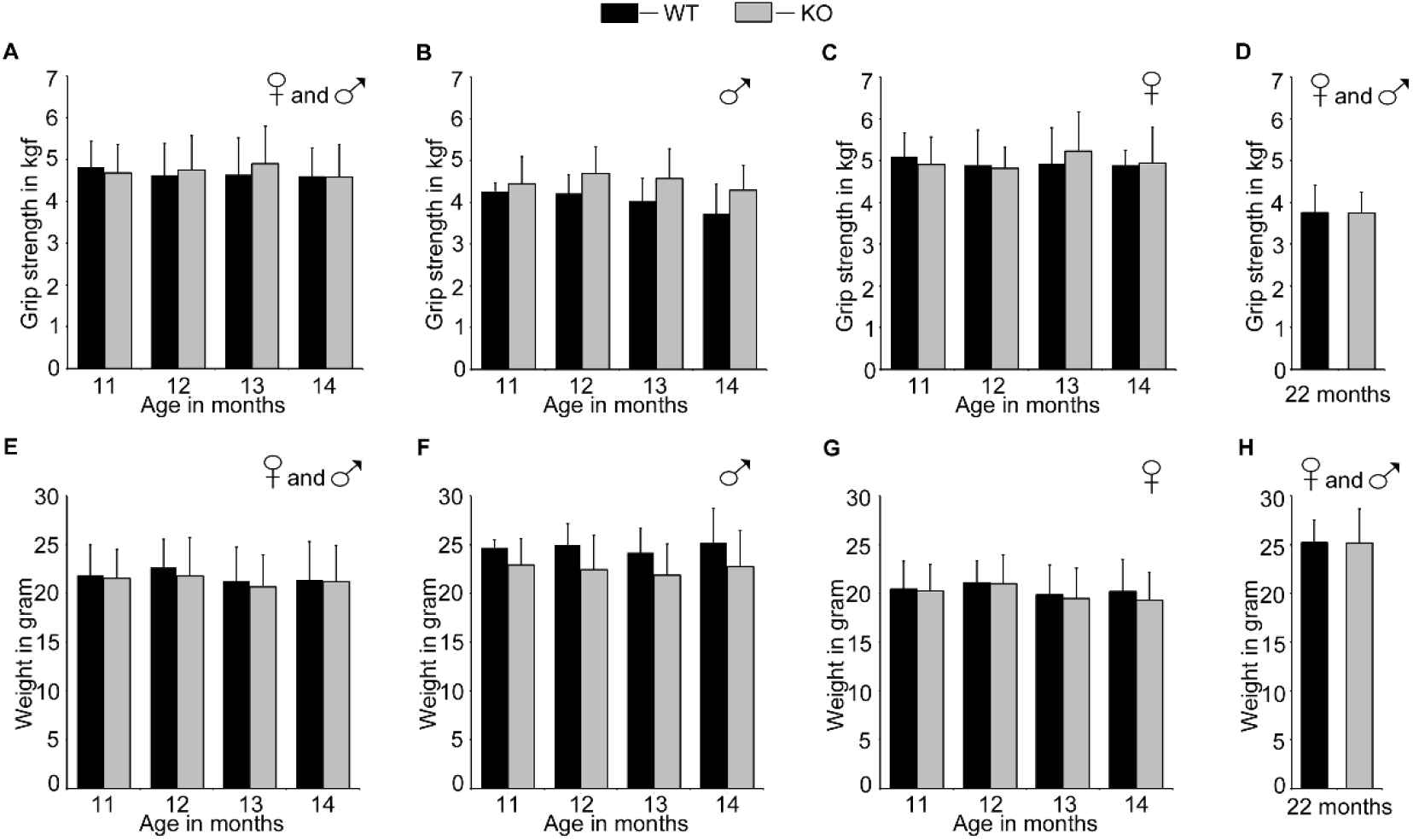
Paw grip strength analysis of optineurin knockout mice: **(A)** Bar diagram showing paw grip strength analysis of WT and optineurin KO mice at the age of 11 (15 WT, 23 KO; p=0.544), 12 (27 WT, 32 KO; p=0.445), 13 (25 WT, 41 KO; p=0.236) and 14 (21 WT, 29 KO; p=0.902) months. **(B)** Bar diagram showing paw grip strength analysis of WT and optineurin KO male mice at the age of 11 (5 WT, 11 KO; p=0.567), 12 (11 WT, 17 KO; p=0.038), 13 (8 WT, 20 KO; p=0.065) and 14 (5 WT, 16 KO; p=0.093) months. **(C)** Bar diagram showing paw grip strength analysis of WT and optineurin KO female mice at the age of 11 (10 WT, 12 KO; p= 0.485), 12 (16 WT, 15 KO; p=0.76), 13 (17 WT, 21 KO; p=0.313) and 14 (16 WT, 13 KO; p=0.8) months. **(D)** Bar diagram showing paw grip strength analysis of WT and optineurin KO mice at the age of 22 months (7 WT, 17 KO; p=0.95). **(E)** Bar diagram showing body weight analysis of WT and optineurin KO mice used in **(A)** at the age of 11 (p= 0.741), 12 (p= 0.269), 13 (p=0.488) and 14 (p=0.872) months. **(F)** Bar diagram showing body weight analysis of WT and optineurin KO male mice used in **(B)** at the age of 11 (p=0.179), 12 (p=0.047), 13 (p=0.082) and 14 (p=0.208) months. **(G)** Bar diagram showing body weight analysis of WT and optineurin KO female mice used in **(C)** at the age of 11 (p=0.875), 12 (p=0.911), 13 (p=0.691) and 14 (p=0.454) months. **(H)** Bar diagram showing body weight analysis of WT and optineurin KO mice used in **(D)** at the age of 22 months (p=0.99). Quantitation represents mean ± SD.

Wire hanging experiment is a method used to measure muscle strength, coordination and endurance in mice. We designed a modified rotating wire apparatus based on a previously described model by which one measures the hanging time of mice [16]. Mice were suspended on a freely rotating 50 inches long wire welded into a circle. This wire was suspended on a pulley which allowed the wire to rotate on application of downward force by the mouse. The mouse has a tendency to move on the wire by pulling it. For pulling the wire and moving on it, muscle coordination plays an important role. Thus, this method gives an estimate of muscle strength, coordination and endurance in mice. Sixteen month old WT control and Optn KO mice were suspended on the rotating wire and the duration of hanging on the wire was recorded. It was observed that though the Optn KO mice showed slightly shorter hanging time as compared to the control WT mice, the difference was not significant (Figure 3A). The normalized hanging strength is calculated by multiplying the hanging time in seconds by the weight of the animal in grams. We observed that KO mice showed lower normalized hanging strength as compared to control WT mice, but the difference was not significant statistically (Figure 3B).

**Figure 3:**
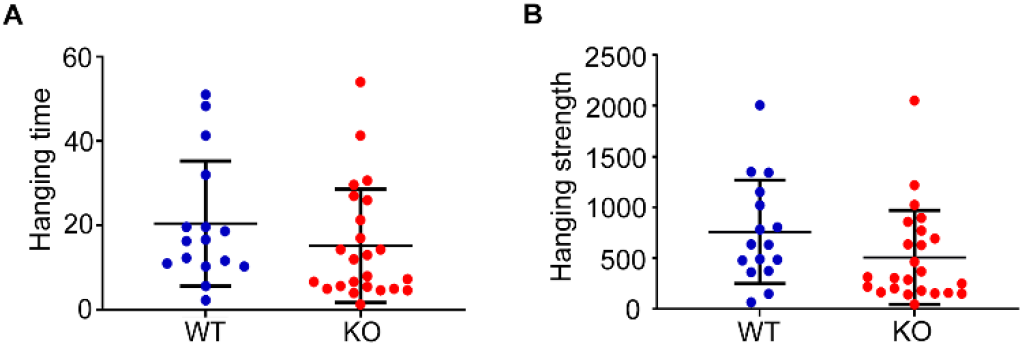
Muscle strength analysis of optineurin knockout mice by rolling wire apparatus: Rolling wire test was performed with 16 months optineurin KO (24) and WT (16) mice. **(A)** Graph showing the time in seconds spent by mice hanging on the rolling wire (p=0.25). **(B)** Graph showing the normalized hanging strength calculated by multiplying the hanging time in seconds by the weight of animal in grams (p= 0.11). Quantitation represents mean ± SD.

## Discussion

Several ALS-associated mutations of OPTN are deletions or truncations, which are likely to cause loss of function of OPTN. Therefore, the loss of Optn in mice is expected to cause ALS. However, we observed that Optn-deficient mice did not develop features of ALS, such as bending of back, loss of body weight, decrease in paw grip strength even at 11-22 months of age. Therefore, we suggest that the loss of optineurin alone is not sufficient to cause ALS in mice. Loss of optineurin may require complementation with some other genetic or environmental factors for manifestation of neuro-degenerative phenotypes associated with ALS. However, we observed that a significant percentage of *Optn^-/-^* mice show patchy hair loss. Patchy hair loss is a common symptom of a clinical condition known as alopecia. Alopecia areata is an autoimmune disease, which is caused by damage of hair follicles due to activation of T cells [20]. It has been reported that certain subgroups of cytotoxic T cells (CD8 ^+^ T cells), are both necessary and sufficient for the induction of alopecia areata [20, 21]. Further work is required to understand the molecular mechanisms underlying hair loss caused by deficiency of optineurin.

Ito et al, generated C57BL/6 *Optn^-/-^* mice by Cre-Lox recombination system and used them to investigate the mechanism by which loss of optineurin could lead to ALS [22]. They observed that the number and morphology of motor neurons in spinal cord of *Optn^-/-^* and WT mice were similar. However, they observed reduction in the number of motor axons and abnormal myelination in the ventrolateral white matter of spinal cord in *Optn^-/-^* mice. They also observed denervation of neuromuscular junctions in the tibialis anterior muscle in *Optn^-/-^* mice. They suggested that deficiency of optineurin leads to axonal pathology without affecting the motor neuron cell bodies [22]. They further suggested that optineurin deficiency specifically in the oligodendrocytes and myeloid cells promote axonal loss and dysmyelination in the spinal cords of *Optn^-/-^* mice. They did not observe any defect in locomotor activity in *Optn^-/-^* mice, as analyzed by open field test. They reported that vertical rearing activity was significantly reduced in *Optn^-/-^* mice and hence suggested that optineurin deficiency leads to hind-limb weakness. However, they had not analyzed the front paw grip strength of *Optn^-/-^* mice by a paw grip strength meter.

Munitic et al. had generated a mouse model, Optn^470T^, in which the entire ubiquitin binding domain in optineurin was deleted [23]. They observed that in 129 X C57BL/6 Optn^470T/470T^ mice, there was embryonic lethality and incomplete penetrance. However, upon backcrossing to C57BL/6, the viability of offsprings was restored. This observation indicates that background of mice has a significant role in manifestation of phenotype of optineurin deficiency. They also observed that the mice which survived looked similar to wild type mice and had normal distribution of immune cells. It was also reported later that the microglia from Optn^470T/470T^ mice showed reduced TBK1 activation and interferon (IFN)-β production upon stimulation of Toll-like receptor [24]. The authors further suggested that disruption of optineurin/TBK1-mediated IFN-β axis might lead to loss of neuroprotection and this could predispose to neurodegeneration.

Slowicka et al. (2016), had generated C57BL/6 background optineurin knockout mice by deleting exon 3 of optineurin gene using Cre-Lox recombination system [25]. They observed that these mice were born with normal Mendelian segregation and reached adulthood without any developmental abnormality. They observed that optineurin knockout mice, up to 50 weeks’ age, did not develop any phenotypic abnormality. They also observed that the brain of these mice did not show any damage, inflammation or presence of protein aggregates. Overall, the mice did not show any spontaneous inflammatory phenotype. Upon stimulation, there was decreased IRF3 signaling and type I IFN production by bone marrow derived macrophages (BMDMs). Also, these Optn KO mice were more susceptible to *Salmonella* infection [25]. Another group has investigated the effect of optineurin deletion on inflammation and cytokine secretion [26]. They observed that upon bacterial challenge, the BMDMs derived from *Optn^-/-^* mice showed reduced pro-inflammatory cytokine (tumor necrosis factor- α and interleukin 6) production and elevated interleukin 10 and CXCL1 production. They also reported that optineurin provides protection against bacterial infection in mice and that optineurin deficiency results in increased susceptibility to *Citrobacter*-induced colitis.

Another mouse model to understand the role of optineurin in inflammation is the *Optn*Δ^157^ model in which the TBK1-interacting domain in the N-terminus of optineurin is deleted [27]. It was observed that *Optn*Δ^157^ mice were born with normal Mendelian ratio and did not show any signs of disease. BMDMs derived from OptnΔ^157^ had reduced TBK1 activity and mounted low IFN response. The authors suggested that optineurin is a positive regulator of TBK1. Recently, another study has described the role of optineurin in osteoclast differentiation using global optineurin knockout mice generated by Cre-Lox recombination system in C57BL/6 background [28]. Genome wide association studies had previously identified OPTN as a candidate gene for susceptibility to PDB (Paget’s disease of bone) [29]. Wong et al. [28] reported that all the *Optn^-/-^* mice showed clinical features observed in PDB. They reported that ex vivo primary osteoclast differentiation led to increased osteoclastogenesis due to deficiency of optineurin. They suggested that defect in IFN signaling due to deficiency of optineurin could be a causative factor for development of PDB symptoms in *Optn^-/-^* mice.

Recently, it has been shown that Optn-deficient mouse embryonic fibroblasts and mouse tissues show enhanced ER stress response signaling, as shown by increased expression of various ER stress induced genes [30]. The level of IRE1 and PERK proteins, which are sensors and signal transducers of ER stress, was higher in Optn-deficient cells, which possibly leads to enhanced ER stress response signaling and cell death. Because ER stress plays an important role in ALS pathogenesis [31], and Optn modulates ER stress response signaling, Optn deficiency may partly contribute to ALS pathogenesis by potentiating ER stress response signaling.

Thus, Optn deficiency leads to defects in various cells and tissues, particularly in immune cell signaling. Defective signaling in brain cells (neurons, microglia) might contribute to ALS pathogenesis. Our results suggest that Optn deficiency alone is not sufficient to cause ALS. Our results also suggest that Optn deficiency contributes to patchy hair loss, a disorder known to be caused by impaired immune cell function.

## Acknowledgements

GS acknowledges the J.C. Bose National Fellowship grant (SR/S2/JCB-41/2010) from the Department of Science and Technology, Government of India. SCM acknowledges the Council for Scientific and Industrial Research, India for research fellowship. The authors acknowledge Bedaballi Dey, Gopalakrishna Ramachandran and Rajashree Ramaswamy for their help in some experiments.

